# DeepSeqPanII: an interpretable recurrent neural network model with attention mechanism for peptide-HLA class II binding prediction

**DOI:** 10.1101/817502

**Authors:** Zhonghao Liu, Jing Jin, Yuxin Cui, Zheng Xiong, Alireza Nasiri, Yong Zhao, Jianjun Hu

**Affiliations:** Z.Liu was with the Department of Computer Science and Engineering, University of South Carolina, Columbia, SC, 29201; J. Jin, Y. Cui, Z. Xiong,A. Nasiri, Y. Zhao, and J. Hu are with the Department of Computer Science and Engineering, University of South Carolina, Columbia, SC, 29201

**Keywords:** MHC, HLA, Deep Learning, Attention mechanism, Binding core

## Abstract

Human leukocyte antigen (HLA) complex molecules play an essential role in immune interactions by presenting peptides on the cell surface to T cells. With significant progress in deep learning, a series of neural network based models have been proposed and demonstrated with their good performances for peptide-HLA class I binding prediction. However, there still lack effective binding prediction models for HLA class II protein binding with peptides due to its inherent challenges. In this work, we present a novel sequence-based pan-specific neural network structure, DeepSeaPanII, for peptide-HLA class II binding prediction. Compared with existing pan-specific models, our model is an end-to-end neural network model without the need for pre- or post-processing on input samples. Besides state-of-the-art peformance in binding affinity prediction, DeepSeqPanII can also extract biological insight on the binding mechanism over the peptide and HLA sequences by its attention mechanism based binding core prediction capability. The leave-one-allele-out cross validation and benchmark evaluation results show that our proposed network model achieved state-of-the-art performance in HLA-II peptide binding. The source code and trained models are freely available at https://github.com/pcpLiu/DeepSeqPanII.

## 1 Introduction

The human leukocyte antigen (HLA) complex is responsible for presenting peptides on cell surfaces for recognition by T-cells. There are two major groups of HLAs: class I and class II. HLAs class I present peptides that originate from the cytoplasm while HLAs class II present those originating extracellularly from foreign bodies such as bacteria. Another difference between these two classes is how they are expressed. The class I HLA protein has one chain (*α*) and the class II HLA protein has two chains (*α* and *β*). HLA genes are highly polymorphic, which allows them to fine-tune the adaptive immune system. Prediction of HLA-II binding peptides is important to vaccine design and targeted therapy development in immunology and cancer immunotherapy, but is challenging because HLA-II are highly polymorphic and the size of the peptides presented varies [1], [2]. As experimentally characterizing the binding specificity for all HLA molecules is costly in terms of time and labor, effective computational prediction methods are needed for HLA-II peptide binding affinity prediction. It has been shown that computational tools for predicting neoantigens are placing an increasingly important role in interrogating cancer immunity [3].

In the past few years, deep neural networks have achieved great success in computer vision, pattern recognition, and natural language processing [4]. Inspired by that, a series of deep neural network models have been proposed to address the peptide-HLA class I binding problem [5], [6], [7]. In our previous work, we developed a novel deep convolutional neural network based pan-specific model for MHC-I peptide binding prediction [8]. However, for HLA class II, there are few deep neural network based prediction algorithms in the published literture. In 2018, Nielsen et al. established an automated benchmarking platform for MHC class II binding prediction methods [9] while a few benchmark studies have been conducted for MHC class I binding [10], [11], [12]. Currently, NN-align(2009) [13], NetMHCIIpan-3.1(2015) [14], Comblib matrices(2008) [15], SMM-align(2007) [16], Tepitope(1999) [17] and Consensus IEDB (2008) [18] are included in this benchmark study, most of which were developed quite a while ago. More recently, there are several major reports on HLA-II peptide binding prediction [1], [2], [19]. First Garde et al. [2] proposed to take advantage of a large set of MHC class II eluted ligands generated by mass spectrometry to guide the prediction of MHC class II antigen. Next in *Nature Biotechnology*, Racle et al. [19] proposed to combine unbiased mass spectrometry with a motif deconvolution algorithm to analyze peptides eluted from HLA-II molecules. They developed a probabilistic framework to learn multiple motifs on the peptides, as well as the weights and binding core offsets of these motifs. Their probablistic predictor of HLA-II ligands (MixMHC2pred) was shown to outperform the NetMHCIIPanII. At the same time, also in *Nature Biotechnology*, a long short-term memory (LSTM) based recurent neural network model,MARIA [1], was proposed to handle the high variability in the length of HLA-II peptide ligands (826 amino acids). When trained with both traditional HLA binding affinity data, the MS-based antigen presentation profiling datasets together with gene expression data and flanking residues of peptides, their deep learning model achieved significantly better prediction performance. While their focus is on demonstrating the importance of new multi-modal data sources such as peptide HLA ligand sequences identified by mass spectrometry, expression levels of antigen genes and protease cleavage signatures, their deep neural network model is a basic LSTM model with one-hot encoding and two dense layers.

In this paper, we are interested in developing more advanced deep neural network architecture for achieving interpretable Class-II HLA peptide binding affinity prediction and binding core prediction. Compared with HLA class I peptide binding prediction, binding prediction of peptide-HLA class II is much more challenging due to two main facts [20]:

1. **Two amino acid chain structure and highly variable lengths.** HLAs of class I have one protein sequence and all HLA protein sequences have the same length. While in class II, HLA proteins have two amino acid sequences and their sequence lengths vary for different alleles, which causes issues for pan-specific binding prediction methods [8].
2. **Longer peptides.** HLA class I molecules have close-end binding groove. Thus, MHC-I binding peptides are 811 consecutive residues among which 9 peptide nonamers are most common. On the other hand, the groove of MHC-II molecules has open ends, which generally bind to longer peptides, normally 1418 residues. In those long peptides, a small part (usually nine amino acid residues, called binding core) is fitted into the groove, with remaining peptide termini on both ends extending outside [21].

In previous work, several strategies have been proposed to address these two challenges in MHC-II binding prediction. SMM-align, an allele-specific method (each allele has a trained model), uses Metropolis Monte Carlo procedure to search an optimal weight matrix which could be used to calculate the binding affinity given any 9-length peptide [16]. NN-align instead identifies the binding core given a peptide using Gibbs sampling, and then uses this binding core and binding affinity to update network weight [13]. NetMHCIIpan-3.1 is a pan-specific method (one model for all alleles), in which protein sequences are represented as pseudo sequences which were extracted from known binding structures. And then a peptide is processed through SMM-align to generate one binding core peptide and suboptimal peptides (as shown in Figure S1.). All these methods use some kind of alignment or pre-processing on peptides to obtain the binding core of the peptide to overcome highly variable length of the peptides issue. In NetMHCIIpan-3.1, the only pan-specific method, pseudo sequences are used to address variable lengths of different HLA class II proteins. In the latest MARIA algorithm, the variable lenght of peptides are handled with the LSTM layer.

To address two issues seamlessly, here we propose a novel pan-specific deep neural network model with the attention mechanism, DeepSeqPanII (as shown in Figure 1) for MHC-II peptide binding prediction. In our recurrent neural network module, raw peptide and HLA sequences are directly encoded as three vectors of unified-sizes. We then feed these three vectors into a convolutional network to extract binding context information,which is then used to predict binding affinity. A major advantage of our model compared to MARIA and other conventional machine learning models is that by taking advantage of the attention mechanism [22], we could identify the binding core of the peptide based on the attention vector automatically learned by the model during the end-to-end training without any supervised or alignment information. Our contributions in this work are summarized as follows:

**Fig. 1.**
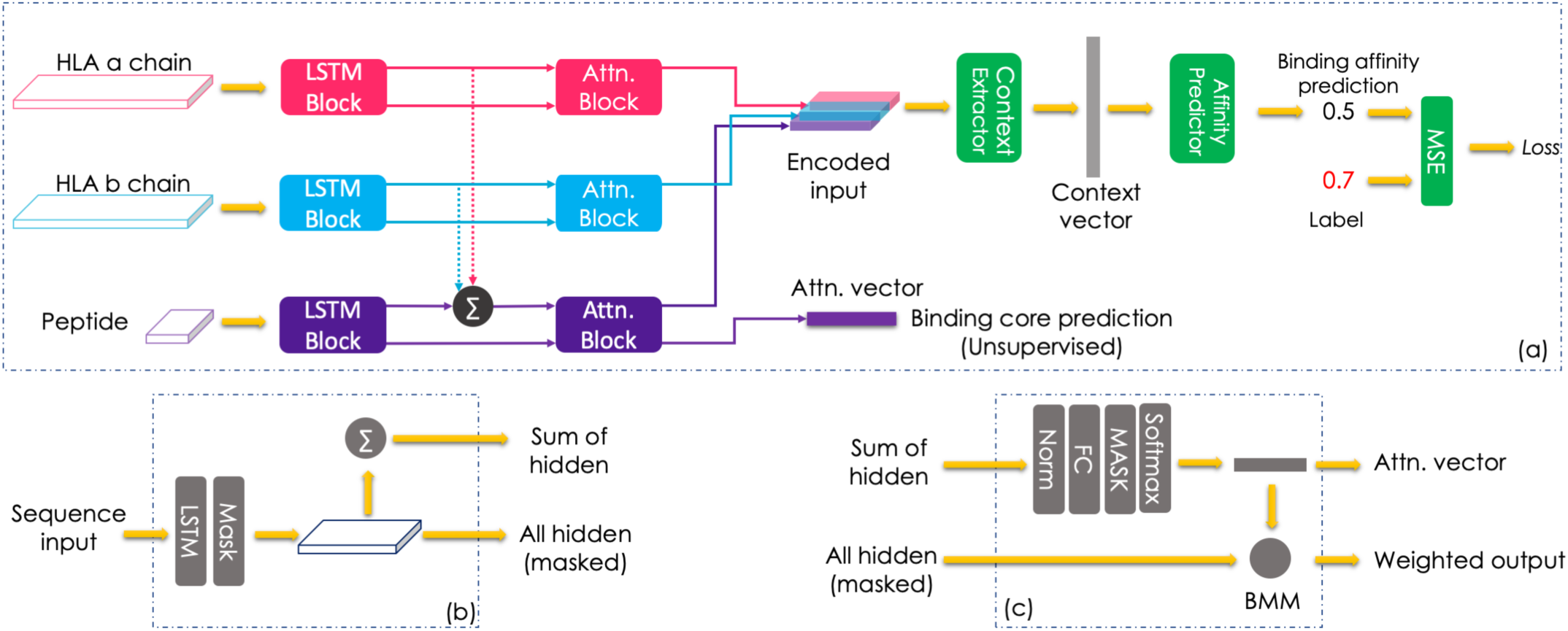
Model architecture of DeepSeqPanII. (a) The overall network structure. (b) Inner structure of LSTM Block. (c) Inner structure of Attention Block.

- We proposed an end-to-end pan-specific deep neural network architecture with attention mechanism for MHC-II peptide binding prediction, in which only raw sequences are needed to train the prediction models.
- We develop a way to interpret the learned weights of our model to identify the peptide binding core ab initio, which demonstrates that our model could learn insightful knowledge of the binding mechanism in an unsupervised way.
- Based on extensive benchmark experiments, we showed that our model could achieve state-of-the-art prediction performance.
- The DNN architecture, source code and pre-trained models are all available for downloading for others to reproduce our work.

## 2 Materials and methods

### 2.1 Dataset

In this work, we collected two training data sets: BD2013 and BD2016 which were generated in 2013 [23] and 2016 [24], respectively. Both data sets were sourced from the IEDB ^1^. In the related works, NetMHCIIPan 3.0 was trained on BD2013 and NetMHCIIPan 3.2 was trained on BD2016. Recently, Andreatta et al. setup an automated platform to benchmark peptide-MHC class II binding prediction methods [9]. We downloaded all available benchmark datasets from 2016-12-31 to 2017-12-29 and evaluated our model on this dataset.

HLA class II protein sequences were downloaded from IPD-IMGT/HLA [25] database. With downloaded *α* sequences and *β* sequences, we performed two multiple-sequence alignments separately. Online alignment tool Clustal Omega ^2^ supported by EMBL-EBI [26] was used here,which is a fast multiple-sequence alignment tool that only requires a list of protein sequences as input. All datasets used in our experiments are included in our GitHub repository for this project.

### 2.2 Sequence encoding

We mix one-hot encoding and BLOSUM62 matrix to represent a protein sequence. Given a sequence with *L* amino acids, it is encoded as a 2D tensor with dimension *L*(*length*) × 43(*channel*). We tried several permutations of one-hot, BLOSUM62 and physical properties. At last, we found this combination gave us best performance. To reduce the training time, we padded all sequences to maximum lengths such that they can be processed in batch during the training stage. More specifically, we set the encoding lengths of peptide sequences, HLA *α* and *β* sequences as 25, 274 and 291 respectively. Theese values were obtained by identifying the maximum lengths for all the HLA *α* and *β* sequences and peptides in our training dataset. Though we padded our input sequences during training, in evaluation stage, LSTM encoders can accept input of arbitrary length.

### 2.3 LSTM-CNN model with Attention mechanism

Figure 1 shows the neural network architecture of our DeepSeaPanII model. It is developed based on our previous work DeepSeqPan for MHC-I binding prediction, which includes a peptide encoder, a HLA encoder, a binding context extractor, and binding affinity predictor. Here, our DeepSeqPanII network also has three parts but with different configuration compared to DeepSeqPanI: i) sequence encoders, ii) binding context extractor and iii) affinity predictor. Given a sample of HLA *α* chain, HLA *β* chain and peptide, sequence encoders encode them as shape-unified output tensors. These three encoded tensors are expected to extract key features that contribute to the binding. Binding context extractor will then take three encoded tensors and output a vector encoding the binding context between this allele and the peptide. It will learn the coupling relationship between the peptide and the HLA sequence. Finally, based on this binding context vector, the predictor will be trained to predict the binding affinity.

As we discussed above, unlike HLA class I, a HLA class II receptor consists of two amino acid sequences with highly variable lengths. Also, since HLA class II binding pockets are open pockets, the peptide sequence lengths range from 8 to 26 in general. To address these two issues, choosing proper encoder architecture becomes critical. Compared to the encoder design, We found that the structures of the context extractor and the affinity predictor do not have dramatic impact on DeepSeqPanII’s prediction performance. At first, we tried to use convolutional neural networks as encoders as we did in DeepSeqPan. However, we could not achieve satisfactory performance with this encoder architecture. A possible reason is that the lengthsof protein and peptide sequences of class II are highly variable. In DeepSeqPan for HLA class I, since all protein and peptide sequences have the same lengths, this was not a problem. To address this highly variable length issue, we propose to use LSTM for encoding the peptide and HLA sequences, which has obtained great success in natural language processing. Also, considering the existence of binding motifs in both the HLA alpha and beta sequences and the peptides, we added an attention module to each of the three encoders [27] in our model to learn the importance of different positions to peptide binding.

Below, we introduce details of each part in the DeepSeqPanII model as illustrated in Figure 1:

a. **DeepSeqPanII network.** The overall network is composed of three sequence encoders, a binding context extractor, and an affinity predictor. An input sample consists of three parts: HLA *α* chain, *β* chain and a peptide. Three sequences are encoded as discussed in previous section. For each sequence, it will be fed into its LSTM block (*LSTM Block* in Figure 1(a)) first. The LSTM block will output two tensors: hidden state tensor and sum of all hidden state tensor. Then these two tensors go into the attention block (*Attn. Block* in Figure 1(a)), which will compute weighted output and the associated attention vector. For attention vectors, they are not directly used to predict final binding affinity values. However, the attention vectors could give us insight over the position importance to the final binding affinity prediction. With three weighted output obtained from the attention blocks, we combine them along channel axis as a new tensor (*Encoded input* in Figure 1(a)), which will be fed into the convolutional network (*Context Extractor* in Figure 1(a)). This convolutional network then outputs a 1D vector (*Context vector* in Figure 1(a)), which encodes all binding context information of this sample. It will go through a fully connected network (*Affinity Predictor* in Figure 1(a)) to calculate the final predicted binding affinity value.
b. **LSTM block.** The input of the LSTM block is the encoded sequence tensor with dimension *L*(*length*) × *C*(*channels*) and the sequence mask vector with dimension *L*(*length*). The encoded sequence tensor is fed into the LSTM layer with *N* hidden states. The LSTM layer will output a tensor with dimension *L × N*. With this raw output tensor and the sequence mask vector, we manually masked the raw output by assigning padding positions’ output values as 0. The reason for masking output is that even though LSTM can handle variable-length input, we padded all input sequences to the same length *L* for easier training of the deep neural network. It allows batch training and speeded training, which is a common setup in LSTM training. By output masking, it could make the network learn faster instead of leting itself figure out the padding’s existence via learning. In our experience, we found that masking can also improve network performance. After output masking, a masked tensor with dimension *L×N* is obtained, which is one of the two output tensors from this block. An additional summarization operation is further applied on this marked tensor to get another output tensor with dimension *N*. This tensor would be used as one of inputs for the attention block.
c. **Attention block.** The attention block aims to extract weighted information based on all hidden states output from the LSTM layer. With the summarization tensor of all hidden states from LSTM as the input signal, it is first normalized and then fed into a fully connected (FC) layer with *L* hidden units. After the FC layer, a vector of *L* size is then calculated, which will be masked again and be fed into the SoftMax layer to calculate an attention vector. After calculating the attention vector, a batch matrix-matrix product (BMM) operation will be applied on the attention vector and all hidden states tensor. The result is a vector with dimension *N*, which contains weighted information of all previous hidden states.

The detailed parameter setup of our network are set as follows.

- **LSTM layer**. The LSTM has 100 hidden units and 2 stacked layers.
- **Context Extractor** The Context Extractor consists of 4 1-D convolutional layers and a max pooling layers.
- **Affinity Predictor** Affinity Predictor is composed of three fully connected layers. First 2 layers have 200 hidden units and followed by a LeakyReLU activation layer and a dropout layer with drop rate of 0.25 The last layer has 1 hidden unit and followed by a Sigmoid activation layer.

### 2.4 Network as math functions

To make it clear to understand our network, we denote the whole network as a series of mathematic functions.

#### Encoding

We use ***S***_*α*_, ***S***_*β*_ and ***S***_*p*_ to denote encoded sequence tensors of the HLA *α*, HLA *β* and the peptide sequences. ***M***_*α*_, ***M***_*β*_ and ***M***_*p*_ are the corresponding mask tensors.

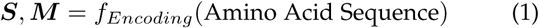

#### LSTM Block

The LSTM block function (Equation 2) takes ***S*** and ***M*** and outputs ***H***_*sum*_ the sum of all hidden states ***H***_*all*_. For peptide, we got 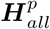 and 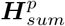. For HLA *α* and *β* chains, we got 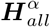, 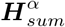, 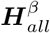 and 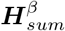.

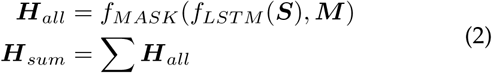

#### Attention Block

The attention block function (Equation 3) for HLA *α* and HLA *β* (Equation) takes ***H***_*all*_ and ***H***_*sum*_ obtained from Equation 2. It outputs an attention vector ***A*** and weighted output ***L***. After attention block, we got ***A***^*α*^, ***L***^*α*^, ***A***^*β*^ and ***L***^*β*^.

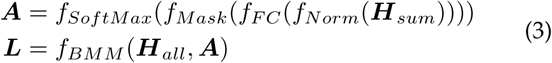

For the peptide sequence, the attention block function (Equation 4) is slightly different since it takes additional inputs: 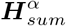 and 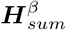. It outputs ***A***^*p*^ and ***L***^*p*^.

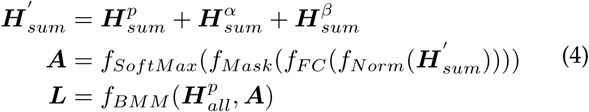

#### Prediction network

After the encoding stage, we have six vectors as output. ***L***_*α*_, ***L***_*β*_ and ***L***_*p*_ are the encoded input vectors for the prediction network together with three attention vectors: ***A***_*α*_, ***A***_*β*_ and ***A***_*p*_. First, we concatenate three encoded input vectors into a new tensor ***L**^I^*′. Then the prediction network function (Equation 5) takes this as input and outputs the binding affinity value *P*.

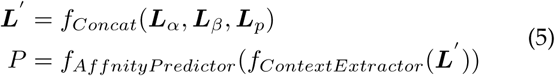

### 2.5 Network training setup

Our batch size is set as 128 and start learning rate is set as 0.01. Our learning rate decay with the *Reduce On Plateau* strategy is adopted here. We reduce the learning rate if the evaluation loss does not decrease for 4 consecutive epochs and we cool down another 4 epochs before checking. The minimum learning rate is 0.0001. We use vanilla SGD as the network optimizer with weight decay regularization (the decay value is 0.01). Also, since training LSTM is sometimes difficult due to gradient exploding, we added gradient clipping into our optimization stage and the threshold is 0.8. Our implementations are based on PyTorch 0.4.1 and all source codes and trained models are freely available at the GitHub repository https://github.com/pcpLiu/DeepSeqPanII.

## 3 Results

### 3.1 Leave one allele out cross-validation

We performed a leave-one-allele-out (LOAO) cross-validation on the BD2016 dataset. We split BD2016 into 54 folds based on allele types. Then, we hold one fold as the testing data and other folds as the training data. The process will be iterated for all folds until we tested on all allele folds. In this way, we mimick the situation where the trained model predicts the binding affinity of unseen allele samples. Actually, one important advantage of pan-specific models with increasing attention over allele-specific models is that they can make predictions on HLA alleles that are not included in the training dataset. For researchers who are interested in alleles without any binding data, this is especially useful.

The LOAO cross-validation results are listed in Table 1. We calculated AUC scores for each validated allele. From the table we observed that out of 54 results, 44 scores of our algorithm are over 0.7, 23 scores are over 0.8 and 3 scores are over 0.9. The results showed that our model has good generalization capability.

**TABLE 1.** LOAO results on BD2016

| Allele | AUC | Allele | AUC |
| --- | --- | --- | --- |
| DQA1*06:01-DQB1*04:02 | 0.694 | DPA1*01:03-DPB1*03:01 | 0.857 |
| DQA1*05:01-DQB1*04:02 | 0.697 | DPA1*01:03-DPB1*02:01 | 0.817 |
| DQA1*05:01-DQB1*03:03 | 0.782 | DRA*01:01-DRB5*01:01 | 0.771 |
| DQA1*05:01-DQB1*03:02 | 0.794 | DRA*01:01-DRB4*01:03 | 0.817 |
| DQA1*05:01-DQB1*03:01 | 0.867 | DRA*01:01-DRB4*01:01 | 0.707 |
| DQA1*05:01-DQB1*02:01 | 0.790 | DRA*01:01-DRB3*03:01 | 0.755 |
| DQA1*04:01-DQB1*04:02 | 0.835 | DRA*01:01-DRB3*02:02 | 0.760 |
| DQA1*03:03-DQB1*04:02 | 0.686 | DRA*01:01-DRB3*01:01 | 0.743 |
| DQA1*03:01-DQB1*03:02 | 0.666 | DRA*01:01-DRB1*16:02 | 0.862 |
| DQA1*03:01-DQB1*03:01 | 0.723 | DRA*01:01-DRB1*15:01 | 0.794 |
| DQA1*02:01-DQB1*04:02 | 0.762 | DRA*01:01-DRB1*13:02 | 0.816 |
| DQA1*02:01-DQB1*03:03 | 0.827 | DRA*01:01-DRB1*13:01 | 0.819 |
| DQA1*02:01-DQB1*03:01 | 0.748 | DRA*01:01-DRB1*12:01 | 0.815 |
| DQA1*02:01-DQB1*02:02 | 0.642 | DRA*01:01-DRB1*11:01 | 0.819 |
| DQA1*01:04-DQB1*05:03 | 0.689 | DRA*01:01-DRB1*10:01 | 0.900 |
| DQA1*01:03-DQB1*06:03 | 0.817 | DRA*01:01-DRB1*09:01 | 0.783 |
| DQA1*01:02-DQB1*06:02 | 0.832 | DRA*01:01-DRB1*08:02 | 0.799 |
| DQA1*01:02-DQB1*05:02 | 0.672 | DRA*01:01-DRB1*08:01 | 0.811 |
| DQA1*01:02-DQB1*05:01 | 0.797 | DRA*01:01-DRB1*07:01 | 0.840 |
| DQA1*01:01-DQB1*05:01 | 0.833 | DRA*01:01-DRB1*04:05 | 0.763 |
| DPA1*03:01-DPB1*04:02 | 0.865 | DRA*01:01-DRB1*04:04 | 0.795 |
| DPA1*02:01-DPB1*14:01 | 0.900 | DRA*01:01-DRB1*04:03 | 0.770 |
| DPA1*02:01-DPB1*05:01 | 0.877 | DRA*01:01-DRB1*04:02 | 0.667 |
| DPA1*02:01-DPB1*01:01 | 0.857 | DRA*01:01-DRB1*04:01 | 0.751 |
| DPA1*01:03-DPB1*06:01 | 0.988 | DRA*01:01-DRB1*03:01 | 0.647 |
| DPA1*01:03-DPB1*04:02 | 0.759 | DRA*01:01-DRB1*01:03 | 0.651 |
| DPA1*01:03-DPB1*04:01 | 0.872 | DRA*01:01-DRB1*01:01 | 0.790 |

In Figure 2, we compared LOAO results of NetMHCIIpan as included in the original dataset with our results by DeepSeqPanII. We found that on 26 alleles, our method performed better over NetMHCIIpan. If we ignore the records with minimal score difference less than 0.01, our model still performs better than NetMHCIIpan on 19 alleles. Moreover, if we only consider particularly large score difference, there are 3 alleles on which NetMHCI-Ipans scores are at least 0.1 higher than ours. Those alleles are DQA1*02:01-DQB1*02:02, DQA1*05:01-DQB1*04:02 and DRA*01:01-DRB1*04:02. In the opposite, our model has at least 0.1 higher scores than NetMHCIIpan on 4 alleles, those are DQA1*01:02-DQB1*05:01, DQA1*04:01-DQB1*04:02, DQA1*01:01-DQB1*05:01 and DRA*01:01-DRB1*13:02. And our models scores on three of those alleles are higher than the NetMHCIIpans above 0.15. But in most cases (47 of 54), two models delivered very similar performances with less than 0.1 margin (as shown in Figure 2). This indicates that two models have similar performance over most alleles while each one has several groups of alleles that it could achieve more accurate affinity prediction.

**Fig. 2.**
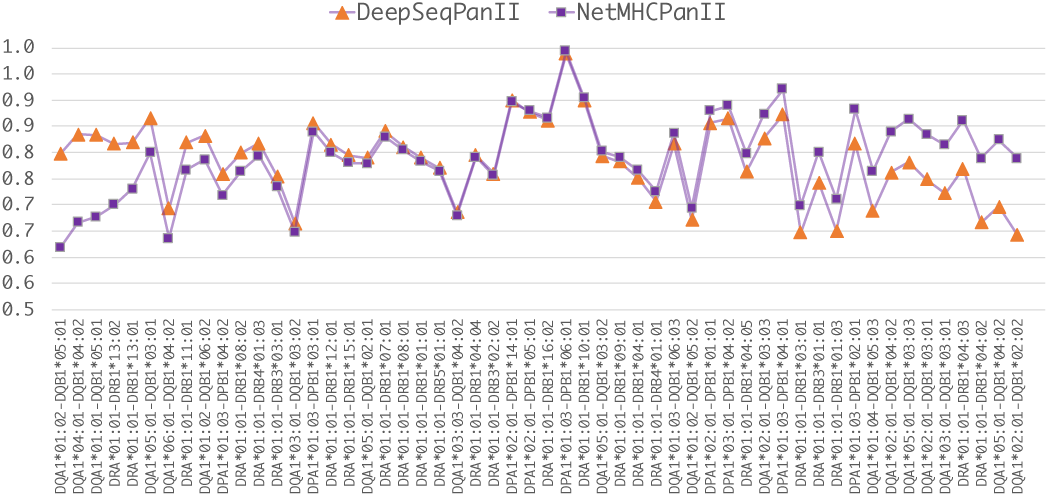
LOAO Performance comparison between DeepSeqPanII and NetMHCIIpan.

### 3.2 Prediction of peptide binding cores via attention mechanism

As we mentioned before, due to the open-ended binding pockets in HLA-II proteins, researchers are usually interested in finding the binding cores of a peptide sequence. In NetMHCIIpan, a pre-processing procedure was used to generate binding core labels. Here, a major goal of our model for binding affinity prediction is to extract insights of the binding mechanism by taking advantage of the attention mechanism, which can learn the importance or contribution of each peptide position to the final binding affinity prediction performance. By inspecting what the attention vectors learned, it makes it possible to predict the binding core of a given peptide sequence. Our intuitive hypothesis is: if the learned neural network pays relatively more attention to some specific amino acid locations of the peptide, these locations may correspond to the binding core.

Here we used a simple approach to determine the binding core from the attention vector: find the subsequence of length-9 with the maximum sum of attention weights (as shown in Figure 3). We conducted testing on the binding core dataset prepared in [14]. In their paper, the authors downloaded the peptide/HLA-DR complexes from PDB database. Then structure-based binding cores were identified by inspecting the location of the bound peptide core within the MHC binding groove. We compared the predicted binding cores on 47 complexes and the results are listed in Table 2. Our of 47 complexes, 8 predicted binding cores match exactly with the experimental ones. In addition, our predicted binding cores missed only one amino acid in 27 complexes. For the remaining 12 complexes, two or more amino acids are missed in our binding core predictions. Overall for about 74% out of the 47 complexex, our attention mechanism based binding core predictions could either exactly match or just miss one amino acids compared to the experimentially determined binding cores. This proves that our deep neural networks with attention mechanisms could capture some key information related to HLA-peptide binding by exploiting large number of training samples.

**TABLE 2.**
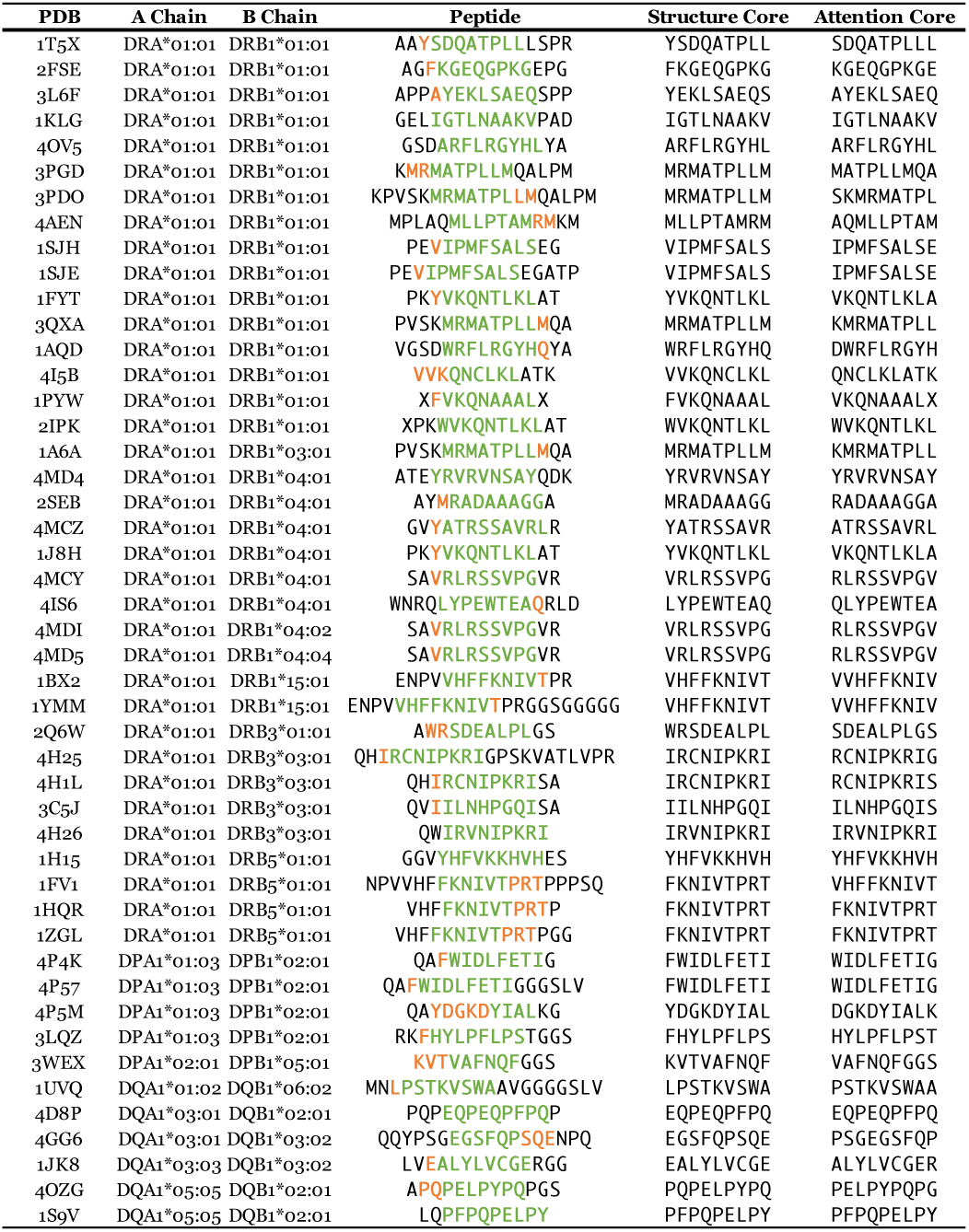
Binding core prediction results. Colored amino acids are structure-based experimental binding cores. Green ones are matched amino acids in our predicted binding cores. Orange ones are missed amino acids in our predicted binding cores.

**Fig. 3.**
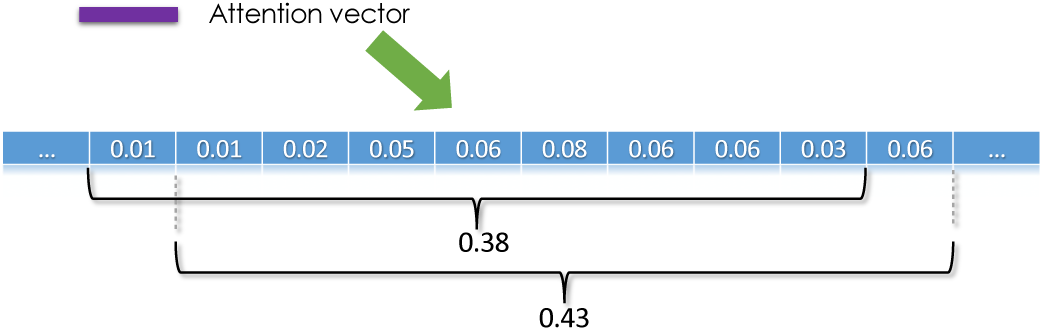
Extract binding core from the peptide attention vector. Using a 9-length sliding window to find out the subsequence with maximum attention weights.

### 3.3 Comparison with other HLA-II binding prediction models on weekly benchmark data

Andreatta et al. recently setup an automated benchmarking for HLA class II alleles [9]. It evaluates participated methods in a similar way as the one for HLA class I [28], which has been widely used to compare performances of different models. We also performed an evaluation for our proposed DeepSeqPanII on the available dataset. Since the benchmark dataset includes data on IEDB since 2014, we trained DeepSeqPanII on BD2013 for a fair comparison. AUC and SRCC scores used in this benchmark were calculated based on all methods’ predictions. The binding prediction results of other methods are included in the original dataset downloaded from the IEDB website. Methods NN-align [13], NetMHCIIpan-3.1 [14], Comblib matrices [15], SMM-align [16], Tepitope [17] and Consensus IEDB [18] are included in this benchmark. We grouped the benchmark data by target alleles and the measurement type. Totally, we have 44 testing groups.

The performance results are listed in Table 3. From the table we can see that, different methods outperformed others on various alleles. And in general, NetMHCIIpan-3.1 and DeepSeqPanII significantly outperform the remaining methods. NetMHCIIpan-3.1 outperforms other methods over 25 test groups in terms of AUC scores and 26 test groups in terms of SRCC scores. DeepSeqPanII outperform all others over 16 test groups in terms of AUC scores and 14 test groups in terms of SRCC scores. NN-align obtains the best AUC scores among 4 groups and best SRCC scores in 3 groups. Surprisingly, Consensus IEDB which combines top performing algorithms does not show a good performance here. One possible reason could be that in the original Consensus IEDB paper, it only included results of several old methods available in 2008. And because their code is not open-sourced, its predictions have not been updated. Since our method and NetMHCIIpan performed overwhelmingly better than other methods on different alleles, an ensemble method based on these two methods could be very promising to improve the overall performance on peptide-HLA class II binding prediction.

**TABLE 3.** Performance comparison on weekly benchmark dataset.

| Allele | Count | Type | DeepSeqPanII |  | NN-align |  | NetMHCIIpan3.1 |  | Comblib |  | SMM-align |  | Tepitope |  | Consensus IEDB |  |
| --- | --- | --- | --- | --- | --- | --- | --- | --- | --- | --- | --- | --- | --- | --- | --- | --- |
|  |  |  | AUC | SRCC | AUC | SRCC | AUC | SRCC | AUC | SRCC | AUC | SRCC | AUC | SRCC | AUC | SRCC |
| HLA-DQA1*01:02/DQB1*05:01 | 825 | IC<sub>50</sub> | 0.60 | 0.21 | - | - | 0.60 | 0.21 | - | - | - | - | - | - | - | - |
| HLA-DQA1*01:02/DQB1*06:02 | 10 | IC<sub>50</sub> | 0.56 | 0.12 | 0.81 | 0.27 | 0.81 | 0.22 | 0.25 | -0.24 | 1.00 | 0.38 | - | - | 0.75 | 0.22 |
| HLA-DQA1*01:03/DQB1*06:03 | 357 | IC<sub>50</sub> | 0.56 | 0.10 | - | - | 0.81 | 0.42 | - | - | - | - | - | - | - | - |
| HLA-DQA1*02:01/DQB1*03:01 | 818 | IC<sub>50</sub> | 0.75 | 0.48 | - | - | 0.81 | 0.59 | - | - | - | - | - | - | - | - |
| HLA-DQA1*02:01/DQB1*03:03 | 759 | IC<sub>50</sub> | 0.76 | 0.50 | - | - | 0.76 | 0.54 | - | - | - | - | - | - | - | - |
| HLA-DQA1*02:01/DQB1*04:02 | 765 | IC<sub>50</sub> | 0.56 | 0.11 | - | - | 0.52 | 0.04 | - | - | - | - | - | - | - | - |
| HLA-DQA1*03:01/DQB1*03:02 | 18 | IC<sub>50</sub> | 0.36 | 0.02 | 0.96 | 0.62 | 0.97 | 0.55 | 0.83 | 0.42 | 0.95 | 0.72 | - | - | 0.99 | 0.70 |
| HLA-DQA1*03:03/DQB1*04:02 | 567 | IC<sub>50</sub> | 0.57 | 0.06 | - | - | 0.48 | -0.08 | - | - | - | - | - | - | - | - |
| HLA-DQA1*05:01/DQB1*03:02 | 834 | IC<sub>50</sub> | 0.74 | 0.48 | - | - | 0.77 | 0.57 | - | - | - | - | - | - | - | - |
| HLA-DQA1*05:01/DQB1*03:03 | 564 | IC<sub>50</sub> | 0.76 | 0.47 | - | - | 0.81 | 0.61 | - | - | - | - | - | - | - | - |
| HLA-DQA1*05:01/DQB1*04:02 | 747 | IC<sub>50</sub> | 0.56 | 0.10 | - | - | 0.58 | 0.14 | - | - | - | - | - | - | - | - |
| HLA-DQA1*06:01/DQB1*04:02 | 565 | IC<sub>50</sub> | 0.52 | 0.00 | - | - | 0.50 | -0.06 | - | - | - | - | - | - | - | - |
| HLA-DRA*01:01/DRB1*01:01 | 69 | Binary | 0.85 | 0.54 | 0.83 | 0.51 | 0.84 | 0.52 | 0.66 | 0.25 | 0.79 | 0.44 | 0.23 | -0.42 | 0.79 | 0.44 |
| HLA-DRA*01:01/DRB1*01:01 | 1070 | IC<sub>50</sub> | 0.81 | 0.63 | 0.78 | 0.60 | 0.80 | 0.63 | 0.70 | 0.43 | 0.74 | 0.51 | 0.28 | -0.46 | 0.75 | 0.53 |
| HLA-DRA*01:01/DRB1*03:01 | 1001 | IC<sub>50</sub> | 0.76 | 0.50 | 0.79 | 0.60 | 0.85 | 0.71 | - | - | 0.78 | 0.58 | 0.25 | -0.52 | 0.80 | 0.61 |
| HLA-DRA*01:01/DRB1*03:01 | 79 | Binary | 0.49 | -0.01 | 0.59 | 0.15 | 0.62 | 0.21 | - | - | 0.62 | 0.20 | 0.43 | -0.11 | 0.60 | 0.17 |
| HLA-DRA*01:01/DRB1*04:01 | 104 | Binary | 0.73 | 0.39 | 0.77 | 0.47 | 0.76 | 0.46 | - | - | 0.69 | 0.33 | 0.22 | -0.48 | 0.69 | 0.34 |
| HLA-DRA*01:01/DRB1*04:01 | 179 | IC<sub>50</sub> | 0.77 | 0.42 | 0.81 | 0.50 | 0.84 | 0.58 | - | - | 0.67 | 0.34 | 0.44 | -0.19 | 0.68 | 0.34 |
| HLA-DRA*01:01/DRB1*04:04 | 861 | IC<sub>50</sub> | 0.85 | 0.67 | 0.80 | 0.59 | 0.86 | 0.71 | - | - | 0.78 | 0.58 | 0.17 | -0.63 | 0.80 | 0.62 |
| HLA-DRA*01:01/DRB1*07:01 | 1071 | IC<sub>50</sub> | 0.86 | 0.70 | 0.87 | 0.74 | 0.88 | 0.76 | 0.79 | 0.55 | 0.86 | 0.71 | 0.19 | -0.61 | 0.86 | 0.72 |
| HLA-DRA*01:01/DRB1*07:01 | 69 | Binary | 0.80 | 0.52 | 0.86 | 0.63 | 0.88 | 0.65 | 0.66 | 0.27 | 0.85 | 0.60 | 0.26 | -0.41 | 0.84 | 0.59 |
| HLA-DRA*01:01/DRB1*08:01 | 889 | IC<sub>50</sub> | 0.82 | 0.64 | - | - | 0.86 | 0.72 | - | - | - | - | 0.20 | -0.59 | 0.80 | 0.59 |
| HLA-DRA*01:01/DRB1*08:02 | 142 | IC<sub>50</sub> | 0.74 | 0.31 | 0.70 | 0.32 | 0.74 | 0.44 | - | - | 0.39 | 0.19 | 0.72 | 0.41 | 0.32 | -0.15 |
| HLA-DRA*01:01/DRB1*09:01 | 873 | IC<sub>50</sub> | 0.84 | 0.59 | 0.85 | 0.67 | 0.87 | 0.70 | 0.60 | 0.16 | 0.79 | 0.57 | - | - | 0.81 | 0.60 |
| HLA-DRA*01:01/DRB1*09:01 | 34 | Binary | 0.66 | 0.28 | 0.84 | 0.59 | 0.84 | 0.58 | 0.62 | 0.22 | 0.80 | 0.51 | - | - | 0.76 | 0.45 |
| HLA-DRA*01:01/DRB1*11:01 | 66 | Binary | 0.64 | 0.23 | 0.72 | 0.37 | 0.75 | 0.42 | - | - | 0.69 | 0.32 | 0.29 | -0.35 | 0.74 | 0.41 |
| HLA-DRA*01:01/DRB1*11:01 | 1006 | IC<sub>50</sub> | 0.84 | 0.68 | 0.87 | 0.75 | 0.89 | 0.78 | - | - | 0.85 | 0.71 | 0.20 | -0.60 | 0.84 | 0.70 |
| HLA-DRA*01:01/DRB1*12:02 | 17 | Binary | 0.84 | 0.59 | - | - | 0.80 | 0.51 | - | - | - | - | - | - | - | - |
| HLA-DRA*01:01/DRB1*13:01 | 18 | Binary | 0.91 | 0.72 | - | - | 0.86 | 0.63 | - | - | - | - | 0.23 | -0.47 | 0.77 | 0.47 |
| HLA-DRA*01:01/DRB1*13:01 | 866 | IC<sub>50</sub> | 0.82 | 0.59 | - | - | 0.77 | 0.53 | - | - | - | - | 0.21 | -0.55 | 0.79 | 0.55 |
| HLA-DRA*01:01/DRB1*13:02 | 134 | IC<sub>50</sub> | 0.71 | 0.14 | 0.91 | 0.61 | 0.90 | 0.62 | - | - | 0.76 | 0.42 | 0.72 | 0.28 | 0.76 | 0.53 |
| HLA-DRA*01:01/DRB1*14:54 | 854 | IC<sub>50</sub> | 0.85 | 0.66 | - | - | 0.89 | 0.71 | - | - | - | - | - | - | - | - |
| HLA-DRA*01:01/DRB1*15:01 | 167 | IC<sub>50</sub> | 0.85 | 0.65 | 0.74 | 0.53 | 0.76 | 0.50 | - | - | 0.51 | 0.06 | 0.64 | 0.18 | 0.47 | 0.02 |
| HLA-DRA*01:01/DRB1*15:01 | 79 | Binary | 0.52 | 0.03 | 0.61 | 0.17 | 0.58 | 0.12 | - | - | 0.45 | -0.08 | 0.50 | 0.01 | 0.52 | 0.03 |
| HLA-DRA*01:01/DRB1*15:02 | 17 | Binary | 0.85 | 0.51 | - | - | 1.00 | 0.74 | - | - | - | - | 0.17 | -0.48 | 0.83 | 0.48 |
| HLA-DRA*01:01/DRB1*15:02 | 18 | IC<sub>50</sub> | 0.93 | 0.69 | - | - | 0.67 | 0.41 | - | - | - | - | 0.44 | -0.39 | 0.54 | 0.38 |
| HLA-DRA*01:01/DRB3*01:01 | 852 | IC<sub>50</sub> | 0.66 | 0.32 | 0.83 | 0.55 | 0.84 | 0.60 | 0.68 | 0.30 | 0.81 | 0.53 | - | - | 0.80 | 0.48 |
| HLA-DRA*01:01/DRB3*02:02 | 771 | IC<sub>50</sub> | 0.70 | 0.38 | - | - | 0.74 | 0.43 | - | - | - | - | - | - | - | - |
| HLA-DRA*01:01/DRB3*03:01 | 854 | IC<sub>50</sub> | 0.73 | 0.45 | - | - | 0.78 | 0.56 | - | - | - | - | - | - | - | - |
| HLA-DRA*01:01/DRB4*01:01 | 14 | IC<sub>50</sub> | 0.63 | 0.38 | 0.60 | 0.40 | 0.80 | 0.52 | 0.73 | 0.55 | 0.73 | 0.46 | - | - | 0.65 | 0.46 |
| HLA-DRA*01:01/DRB4*01:01 | 18 | Binary | 0.75 | 0.39 | 0.74 | 0.37 | 0.72 | 0.35 | 0.49 | -0.01 | 0.55 | 0.08 | - | - | 0.65 | 0.23 |
| HLA-DRA*01:01/DRB4*01:03 | 839 | IC<sub>50</sub> | 0.80 | 0.58 | - | - | 0.79 | 0.54 | - | - | - | - | - | - | - | - |
| HLA-DRA*01:01/DRB5*01:01 | 18 | Binary | 0.97 | 0.77 | 0.92 | 0.68 | 0.96 | 0.75 | - | - | 0.85 | 0.57 | 0.31 | -0.32 | 0.78 | 0.45 |
| HLA-DRA*01:01/DRB5*01:01 | 762 | IC<sub>50</sub> | 0.79 | 0.63 | 0.81 | 0.66 | 0.84 | 0.74 | - | - | 0.78 | 0.61 | 0.26 | -0.53 | 0.77 | 0.61 |
| Outperformance Count |  |  | 16 | 14 | 4 | 3 | 25 | 26 | 0 | 1 | 1 | 2 | 0 | 0 | 1 | 0 |

## 4 CONCLUSION

In this work, we proposed a novel deep neural network with attention mechanism for peptide-HLA class II binding affinity prediction. Compared with existing pan-specific prediction algorithms, our pan-specific model successfully adopted the attention mechanism, which allows us to extract mechanistic insights of HLA-II peptide binding by interpreting the learned attention vector. The leave-one-allele-out results showed what 44 of 54 (80%) alleles’ AUC scores are over 0.7. This indicated that our model has a good generalization capability, which is a major advantage of pan-specific model since it could predict on unseen alleles. In LOAO experiments, our model and NetMHCPanII delivered similar performance on big part of tested alleles, while one method outperformed the other on some specific alleles. In weekly benchmark test, we also found similar trend. We argue that combining DeepSeqPanII with existing models, researchers could achieve even more accurate prediction results. Examining attention vectors learned by our model, we observed that DeepSeqPanII could *see* the important part of a peptide. By listing attention cores and structure cores side by side, we surprisingly found that our model could capture insightful structural information in an unsupervised way. The proposed sequence encoding approaches and the attention mechanism can be applied to other sequence related bioinformatics problems since it only needs sequence information as input. We are looking forward expanding this approach to other sequence related prediction problems.

## Footnotes

1 www.iedb.org

2 www.ebi.ac.uk/Tools/msa/clustalo/

